# Pretraining Improves Prediction of Genomic Datasets Across Species

**DOI:** 10.1101/2025.08.20.671362

**Authors:** Fangrui Huang, Yitong Wang, Janet Song, Ashok Cutkosky

**Affiliations:** Stanford University; Boston University; Harvard University

## Abstract

Recent studies suggest that deep neural network models trained on thousands of human genomic datasets can accurately predict genomic features, including gene expression and chromatin accessibility. However, training these models is computation- and time-intensive, and datasets of comparable size do not exist for most other organisms. Here, we identify modifications to an existing state-of-the-art model that improve model accuracy while reducing training time and computational cost. Using this stream-lined model architecture, we investigate the ability of models pretrained on human genomic datasets to transfer performance to a variety of different tasks. Models pretrained on human data but fine-tuned on genomic datasets from diverse tissues and species achieved significantly higher prediction accuracy while significantly reducing training time compared to models trained from scratch, with Pearson correlation coefficients between experimental results and predictions as high as 0.8. Further, we found that including excessive training tasks decreased model performance and that this compromised performance could be partially but not completely rescued by fine-tuning. Thus, simplifying model architecture, applying pretrained models, and carefully considering the number of training tasks may be effective and economical techniques for building new models across data types, tissues, and species.

## 1 Introduction

Determining how non-coding genomic sequences regulate the expression of nearby genes is critical to understanding how species, cell types, and disease states arise. Unfortunately, identifying the non-coding sequences that act as regulatory elements in a particular cell type or paradigm has proven to be incredibly challenging. A standard empirical approach to identify active non-coding sequences uses DNA accessibility and biochemical marks such as histone modifications or DNA methylation as proxies for functional activity [23]. However, these experiments are expensive, may require samples that are not be easily accessible, and cannot be feasibly performed on all the sequence differences that exist within normal and disease variation in the same species or that have arisen between species across evolutionary time. Thus, there has been great interest in harnessing existing genomic datasets to build machine learning models that can predict features like DNA accessibility and histone modifications directly from DNA sequence alone.

Several recent efforts to use machine learning in genomics have focused on the task of predicting genomic features directly from the underlying DNA sequence. This approach takes inspiration from the recent dramatic successes of deep learning models in natural language processing, and applies similar models to a genomics setting [4, 5, 6, 15, 31]. One challenge in building models that predict genomic features from the underlying DNA sequence is handling long-range interactions between DNA segments that are up to 100s of kilobases (kb) away from each other. Classical neural network building blocks like convolutional layers are not well-suited to such long-range modeling, but newer techniques like self-attention [25] directly addresses this limitation. Self-attention is the core component of modern large language and computer vision models such as GPT [22], and is a natural candidate to improve genomic modeling by more effectively considering long-range interactions.

One recent model that employs self-attention to incorporate long-range sequence information is Enformer [5]. This model combines both convolutional and self-attention blocks to predict 5313 human genomic datasets and 1643 mouse genomic datasets directly from 196kb input DNA sequences. The Enformer model increases information flow by considering sequences that are up to 196kb from the transcription start site and achieves an impressive 0.625 Pearson correlation coefficient between experimental results and predictions. However, the model has 246 million parameters and is extremely memory and compute intensive to train: the published version was trained on a TPU cluster for 192 TPU days with thousands of human and mouse datasets, and numerous additional resources were likely expended to iterate on different model and data choices. This computationally expensive process makes these techniques inaccessible to many researchers, whom may not have the data, monetary resources, or hardware required to train such a model.

We wondered whether it may be possible to (1) reduce the size of the Enformer model such that it can fit into less advanced (and more commonly available) hardware without compromising model accuracy, (2) whether we can utilize existing Enformer-like models to train new models in different paradigms (pretraining/fine-tuning), and (3) what amount of training data is needed for high accuracy.

“Pretraining” refers to a process where a model is first trained on a large amount of data and the weights from this initial training are then used as the starting point for fine-tuning the model on a task of interest. This pretraining/fine-tuning process is already a standard technique in other areas of machine learning (i.e. computer vision and natural language processing), where it has been used to solve image recognition tasks, question answering tasks, and sentence completion tasks [7, 8, 21, 22] using far less data and compute in the fine-tuning stage than would have been required to train a new model from scratch. This strategy has now begun to be applied to machine learning models that predict genomic features [30], but systematic analyses of how best to perform pretraining and whether pretraining can be applied even for datasets that are very different from the initial training data have been limited. As a challenging, but biologically relevant paradigm, we considered whether pretraining on an Enformer-like model built from thousands of genomic datasets in one species (human) can improve accuracy when a new model is trained on only a single GPU using only one or a small number of genomic datasets in another species.

In this paper, we find that a simplified attention-based architecture derived from Enformer can be trained and fine-tuned on readily available academic hardware to high accuracy (Section 3.1). We then use our simplified architecture to examine models pretrained on human datasets and fine-tuned on datasets from disparate species. Strikingly, we find that this pretraining/fine-tuning process significantly increases accuracy across a wide variety of tasks, tissues, and species while reducing computation time compared to models trained from scratch (Sections 3.2, 3.3). Finally, we find a trade-off between generalization and specialization where training on an excessive number of tasks compromises model performance (Section 3.4). The strategies and techniques demonstrated here can help democratize the use of these models across researchers and experimental contexts and should be generalizable to other models, such as Borzoi [16] and Xpresso [3].

## 2 Methods

### 2.1 Overview

We developed a modified version of the Enformer architecture that can be trained efficiently on a single Nvidia V100 GPU with 16GB of RAM. We then pretrained this architecture on a large human genomic dataset from ENCODE for 10 epochs. This pretrained model was then employed on a variety of downstream fine-tuning tasks involving smaller non-human genomic datasets. These fine-tuned models were compared to models trained from-scratch on the smaller non-human genomic datasets.

### 2.2 Training Data

Our pretraining data consists of the human sequences from the training set developed by [12] and modified by [5] for training Enformer. This dataset has 34,021 training examples, each of which consists of an input DNA sequence of length 196,608 sampled from hg38, paired with 5313 label tracks recorded at a 128 base-pair resolution over the middle 114,688 base pairs of the input sequence.

As fine-tuning data, we employed genomic datasets collected from chickens, rhesus macaques, cows, pigs, and mice. These dataset can be further divided into two types of assays: ATAC-seq data and H3K4me3 ChIP-seq data. ATAC-seq identifies regions of open chromatin and is commonly found at non-coding sequences that act as promoters or enhancers. H3K4me3 is a histone mark that is commonly found at promoters and enhancers [24].

The ATAC-seq data was collected in mice, cows, and pigs [9, 14]. These datasets contain 8935 training sequences for each of mouse, cow and pig. For validation, mouse, cow, and pig have 1048, 1087 and 959 sequences respectively Each sequence is paired with 11 ATAC-seq label tracks for mice, 13 for cows, and 13 for pigs. Note that the Pearson correlation coefficient between replicate ATAC-seq datasets is 0.97 *±* 0.044, placing an upper bound on the accuracy of these experiments. Hyperparameters were tuned on the ATAC-seq data and transferred without tuning for the H3K4me3 experiments.

The H3K4me3 ChIP-seq data was collected in mice, chickens, and rhesus macaques [9, 13, 26]. These datasets contain 10583 training and 1134 test sequences for rhesus, 29952 training and 2209 test sequences for mouse, and 4164 training and 441 test sequences for chicken. We subsampled the mouse and rhesus macaque training datasets to use only 4164 sequences in order to keep the training sets balanced between all three species.

All data was preprocessed using the method and code developed by [13]. Briefly, for each organism, we split chromosomes into shorter contiguous regions, and then randomly partitioned these regions into training and validation or test sets using the same data preprocessing pipeline used to generate the training data for Enformer. We then performed a second preprocessing step to identify regions that are homologous to regions in the original Enformer training data. Regions exhibiting such homology were reassigned to the training set. An approximately equal number of regions that did not exhibit such homology were then reassigned out of the training set.

### 2.3 Loss function, evaluation, and training details

All models were trained to predict track values from raw sequence training data using the Poisson negative log-likelihood as the loss function. Performance was then reported using the Pearson correlation coefficient on the validation set for pretraining and ATAC-seq tasks, or the test set for H3K4me3 ChIP-seq tasks. The Pearson correlation coefficient was defined as 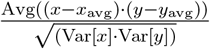 where *x* is the prediction and *y* is the target label value. We used the Poisson negative log-likelihood loss as the optimization target in training instead of directly using the Pearson correlation coefficient because the log-likelihood of a large dataset decomposes as a sum over the log-likelihoods of the individual examples and so is amenable to training using first-order stochastic optimization with small batch size, while the Pearson correlation coefficient is not.

To train the model, we used the AdamW optimizer [17], which is a first-order stochastic optimization algorithm that is a standard choice for deep neural network training. For all experiments, we trained our models for 10 epochs. We tuned the learning rate and weight decay via grid search. For learning rate, we considered values in *{*1e-6, 3e-6, 1e-5, 3e-5, 1e-4, 3e-4*}*. For weight decay, we considered values in *{*1e-4, 3e-4, 1e-3*}*. We performed grid search over the ATAC-seq datasets and recorded the average validation Pearson correlation coefficient for each of the three species (cow, mouse, and pig) and for both fine-tuning and training from scratch. The most common optimal setting was a learning rate of 3e-5 and weight decay of 1e-4. In cases where this was not optimal, the correlation coefficient of the best-performing combination was less than 0.003 larger, which we deemed to be negligible. Therefore, we used this learning rate and weight decay setting for all other tasks with no further tuning. We used a linear learning rate warm-up from 0 to 3e-5 in the first epoch and a linear learning rate decay from 3e-5 to 0 in the later 9 epochs. We used a batch size of 1, as even a batch size of 2 did not fit into GPU memory.

All of our results involve experiments in which models were pretrained on the same human genomic data used to train Enformer (see Methods) and then fine-tuned on one or a few genomic datasets from a non-human species, or in which models were trained from scratch only on datasets from a non-human species with the standard random initialization weights provided by Pytorch (i.e. Kaiming uniform [10]).

All training was performed on a single Nvidia V100 GPU housed at Boston University’s Shared Compute Center. All models were implemented using the Pytorch deep learning framework [20]. The code used for experiments in this paper is available at https://github.com/optimizedlearning/genomicsML.

#### 2.3.1 Single-Track and Multi-Track training

Our ATAC-seq datasets contain many tracks for each species. For these datasets, we trained in two modes: a “multi-track” mode in which the training objective is the average Poisson negative log-likelihood over all tracks and the reported metric is the average Pearson correlation coefficient over all tracks, and a “single-track” mode in which one individual track is selected. The training objective is the Poisson negative log-likelihood for just this selected track and the reported metric is the Pearson correlation coefficient for this same track.

In all cases, our pretrained model was trained for 10 epochs, and the fine-tuned and trained from scratch models were also trained for 10 epochs.

### 2.4 Fine-tuning

To fine-tune a pretrained model on a non-human dataset, we loaded the pretrained weights from a checkpoint file. Then, we deleted the last linear “head” layer that outputs predictions for each of the 5313 human tracks. We replaced this head layer with a new, randomly initialized, head layer that outputs *N* predictions where *N* is the number of tracks in the fine-tuning dataset. Finally, this model is trained on the non-human dataset.

In some experiments, we also froze the pretrained weights. This means that after loading the pretrained model from the checkpoint and replacing the head layer, we disabled gradient computation for all layers in the model except for the head layer. Then, during training, only the weights for the randomly initialized head layer were updated.

### 2.5 Model Architectures

The Enformer model consists of 7 convolutional residual blocks, 11 self-attention blocks, a pointwise convolutional layer and finally two linear “head” layers, corresponding to mouse and human tracks. (see Figure 1) The input to the model is a one-hot encoded DNA sequence of length 196,608 bp where each bp is represented as a 4-dimensional vector: A = [1,0,0,0], C = [0,1,0,0], G = [0,0,1,0], T = [0,0,0,1], N = [0,0,0,0]. For human input data, the output of the model provides predictions for 5313 tracks recorded at a 128 bp resolution over the middle 114,688 bp of the input, corresponding to 114,688/128=896 values per track. For mouse input data, the model provides predictions for 1643 tracks. Thus, the output size for the human head is (896, 5313) and the output size for the mouse head is (896, 1643).

**Figure 1.**
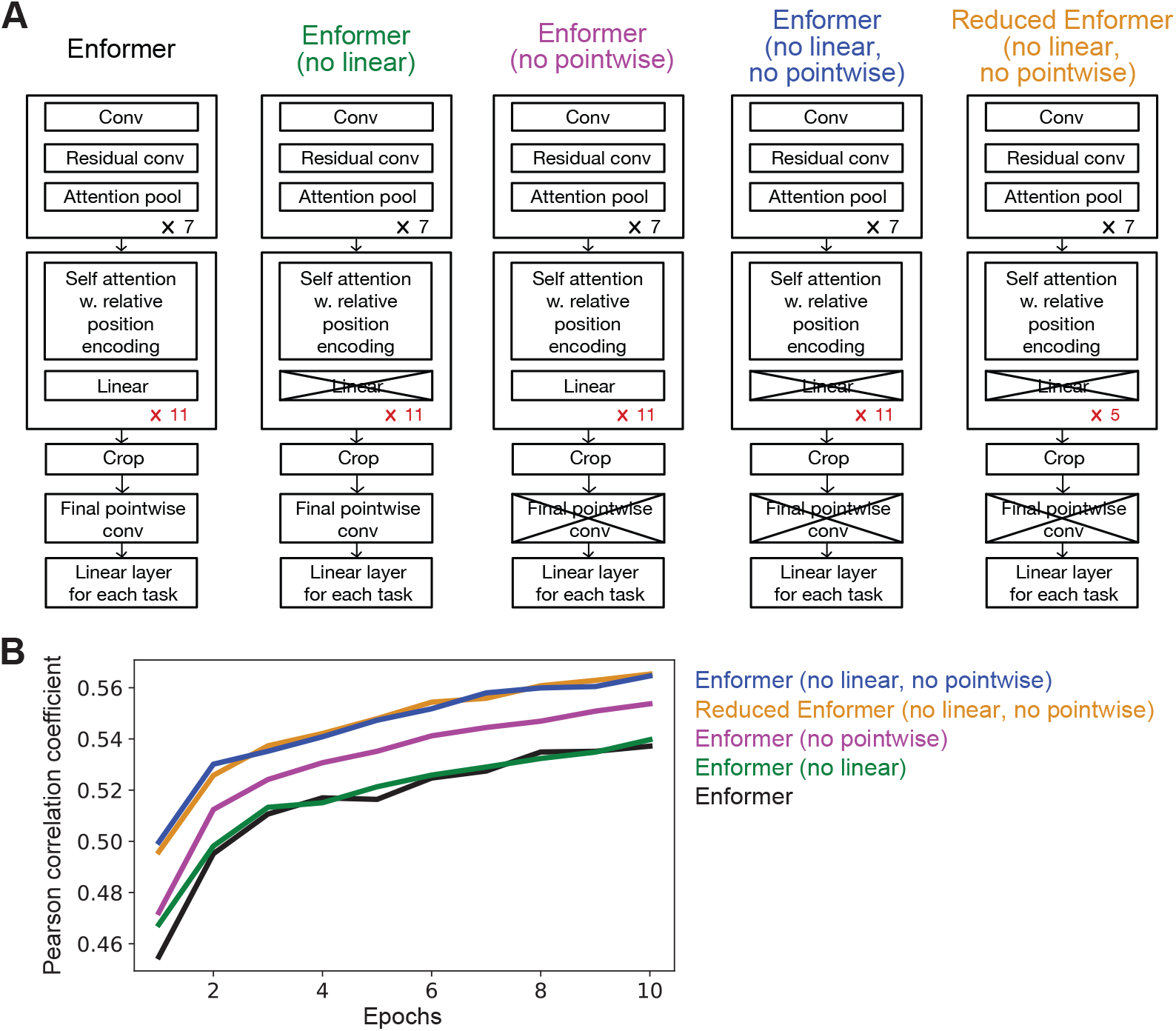
Comparison of different model architectures. (A) We compared model architectures for the pretraining model that had different numbers of attention blocks (11 or 5), were with or without the linear layer in the attention block, and were with or without the final pointwise convolutional layer. (B) Validation Pearson correlation coefficients for different model architectures. Models were trained for 10 epochs on all human datasets that were used to train Enformer. Maximum Pearson correlation coefficient: Enformer = 0.537, Enformer (no linear) = 0.540, Enformer (no pointwise) = 0.554, Reduced Enformer (no linear, no pointwise) = 0.565, Enformer (no linear, no pointwise) = 0.565.

As a baseline, we trained the original Enformer model on the human genomic dataset for 10 epochs. We then trained several variants of the Enformer architecture also for 10 epochs. These variants were: (1) removing the final pointwise convolutional layer, (2) removing a linear layer from the attention blocks, (3) removing both the final pointwise convolutional layer as well as the linear layer from the attention blocks, and (4) reducing the number of attention blocks from 11 to 5.

Of these, we used variant (3) for all downstream analyses.

## 3 Results

### 3.1 Simplifying the Enformer model architecture improves performance in a computationally restricted setting

We first investigated whether we could simplify the Enformer architecture such that it could be easily trained and fine-tuned on cheaper hardware without compromising the model’s performance. We specifically targeted the Nvidia V100 GPU, which is relatively accessible in the academic community. The published Enformer model consists of 7 convolutional residual blocks, 11 self-attention blocks, a pointwise convolutional layer and finally two linear “head” layers, corresponding to human and mouse tracks respectively [5]. We considered several simplifications in which we removed individual layers or reduced the number of repeated blocks in Enformer (Figure 1A).

Enformer was originally trained on both 5313 human datasets and 1643 mouse datasets (see Methods). In order to test whether an Enformer-like model trained in one species can be applied to other species, we limited our training set to just the 5313 human datasets. We also reduced training time (10 epochs vs 150 epochs in the original study) and batch size (1 vs 64) to fit a smaller computational budget. Under these restrictions, the base Enformer model achieved a validation Pearson correlation value of 0.537.

First, we tested whether removing the final point-wise convolutional layer or the linear layer in each self-attention module would affect accuracy (Figure 1B). Removing the final point-wise convolutional layer increased the Pearson correlation coefficient to 0.554 from a baseline of 0.537 (purple in Figure 1B), while removing the linear layer in each self-attention module yielded a marginal increase to 0.540 (green in Figure 1B). Surprisingly, however, removing both the final point-wise convolutional layer and linear layer in each self-attention module further improved the Pearson correlation coefficient to 0.565 (blue in Figure 1B). These changes resulted in a smaller, more efficient model that also achieved better prediction accuracy. By removing the final point-wise convolutional layer and linear layer in each self-attention module the average iteration per second on a V100 GPU increased from 0.755 to 0.765. We used this simplified model architecture for all subsequent experiments.

We also tested whether reducing the number of self-attention blocks in Enformer would affect model accuracy (“Reduced Enformer” in Figure 1). We found that 5 self-attention blocks, rather than the 11 used in Enformer, increased the iterations per second from 0.866 for 11 blocks to 0.92 for 5 blocks, while maintaining a high Pearson correlation coefficient at 0.565. This suggests that there is a low marginal value for additional self-attention blocks at our computational budget. Although we used a model with 11 self-attention blocks in subsequent experiments, reducing the number of self-attention blocks would be a reasonable choice for more resource-constrained researchers.

### 3.2 Pretraining on human datasets improves predictions when fine-tuning on ATAC-seq datasets from non-human species

Using our simplified model architecture, we investigated whether fine-tuning a model pretrained on thousands of human datasets would improve accuracy in predicting ATAC-seq experiments from non-human species when compared to a model trained from scratch without using pretrained weights (see Methods). ATAC-seq identifies regions of open chromatin and is commonly found at non-coding sequences that act as promoters or enhancers [23]. We generated predictions for 11 mouse, 13 cow, and 13 pig ATAC-seq datasets from diverse tissues including muscle, cerebellum, liver, and lung [9, 14] (Figure 2).

**Figure 2.**
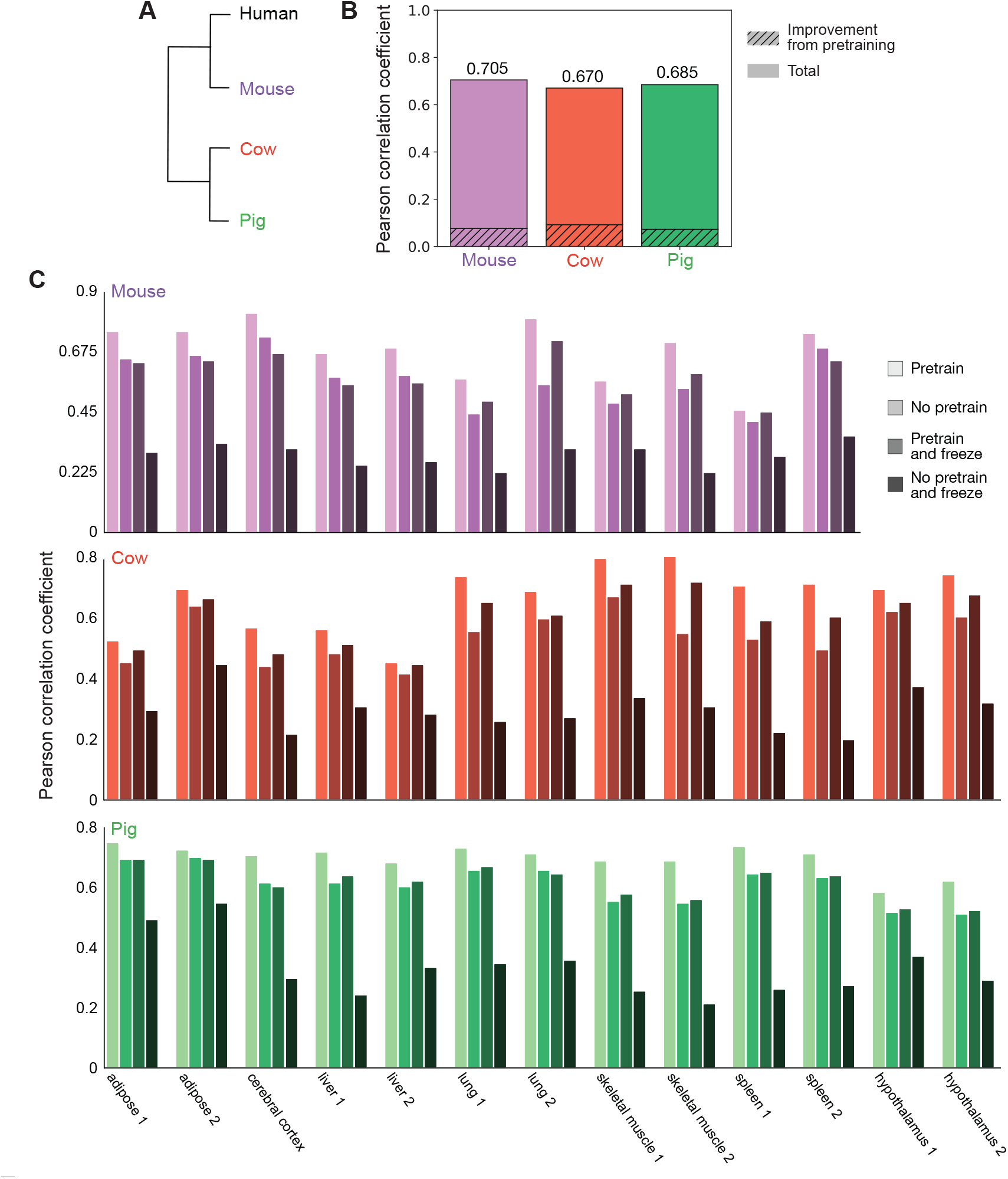
Pretraining improves performance on ATAC-seq datasets across multiple species. (A) Dendrogram of humans and species for which we examined previously generated ATAC-seq datasets (mouse, pig, cow). (B) Validation Pearson correlation coefficients for models trained on 11 mouse, 13 cow, or 13 pig ATAC-seq datasets [9, 14]. The shaded regions are the difference between the Pearson correlation coefficient when using a model pretrained on human genomic datasets that is then fine-tuned on ATAC-seq datasets from mouse, cow, or pig and a model trained from scratch without pretraining. Total height indicates the Pearson correlation coefficient of the fine-tuned model. (C) Validation Pearson correlation coefficients for models trained on individual ATAC-seq datasets from mouse, pig and cow. “Pretrain” is a model fine-tuned from a pretrained model. “No pretrain” is a model trained from scratch. “Pretrain and freeze” is a model fine-tuned from pretrained weights where only the last linear layer is updated in the fine-tuning process. “No pretrain and freeze” is a model trained from scratch where only the last linear layer is updated in the training process.

To investigate whether pretraining improves model performance, we first determined the average Pearson correlation coefficient between experimental results and predictions from the model with and without pretraining for all ATAC-seq datasets for a given species. The average validation Pearson correlation coefficient across tissues is 0.628, 0.5779, and 0.6124 for mouse, cow, and pig respectively without pretraining. Strikingly, these correlation coefficients increased after pretraining to 0.705, 0.670, and 0.685, for an increase of 12.26%, 15.97%, and 11.85% respectively (Figure 2B).

We then asked whether improvements from pretraining persist even when fine-tuning on a *single* ATAC-seq dataset. We compared the Pearson correlation coefficient for models individually trained on the same ATAC-seq datasets from mouse, cow, and pig that were also analyzed above [9, 14].

We examined ATAC-seq datasets for adipose tissue, the cerebral cortex, liver, lung, skeletal muscle, and spleen for all three species. For pig and cow, we also included datasets generated from the hypothalamus. For many of these tissues, replicate datasets were available, and we analyzed these separately.

When training on a single ATAC-seq dataset, pretraining increased the validation Pearson correlation coefficient between experimental results and predictions from the model with an average improvement of 19.24% *±* 10.83% (Figure 2C, compare “Pretrain” to “No pretrain”). This improvement in performance was greater than the improvement observed when training on multiple ATAC-seq datasets. Notably, when averaged across single-dataset experiments, the validation Pearson correlation coefficient after pretraining was 0.691, 0.665, and 0.694 for mouse, cow, and pig respectively, in line with the correlation coefficients achieved when training on multiple ATAC-seq datsets after pretraining. This suggests that pretraining on thousands of human datasets allows for high model accuracy even when fine-tuning on just a single dataset from a non-human species.

Model accuracy varied widely between datasets both across and within species. Pretraining improved the correlation coefficient over training from scratch by 19.83% +/− 11.69% for mice, 23.55% +/− 12.51% for cows, and 14.43% +/− 6.03% for pigs across single-dataset experiments. This is not correlated with evolutionary distance, as mice are more closely related to humans than cows. There was also dramatic variation across tissues and even within replicates of the same tissue within a species. For instance, the Pearson correlation coefficients for spleen 1, spleen 2, and cerebral cortex in mouse were 0.4511, 0.7388, and 0.8114 respectively. This inconsistency in prediction accuracy not only for different species or for different tissues within the same species, but also for replicates of the same tissue in the same species, suggests that differences in model accuracy are due to technical features of the experimental datasets and not due to biological differences between tissue types or species.

Next, we considered freezing the pretrained layers (i.e. not updating the parameters and only updating the last linear layer) on a single ATAC-seq dataset. This is a standard strategy for efficient fine-tuning [29], and if it could achieve comparable effects to “full” fine-tuning, then it would decrease the computational resources on a V100 GPU: the peak CUDA memory usage was 2.49 GB with freezing, compared to 9.50 GB without freezing. We also considered the control group where we perform no pretraining (i.e. random initialization) and update only the last linear layer on a single ATAC-seq dataset (Figure 2C). As expected, only updating the last linear layer without pretraining performed poorly (“No pretrain and freeze” in Figure 2C). Freezing the pretrained layers and only updating the last linear layer on a single ATAC-seq dataset (“Pretrain and freeze” in Figure 2C) dramatically out-performed the control group and performed comparably to training from scratch on a single ATAC-seq dataset. However, it did not perform as well as using a pretrained model and then fine-tuning the entire model (“Pretrain” in Figure 2C).

These experiments show that pretraining followed by fine-tuning the entire model produce the best correlation between experimental and predicted results. However, we note that if computational resources are scarce, using the pretrained model and only updating the last linear layer produces comparable results to updating weights for the whole model without pretraining. While training the whole model took 28h on a Tesla V100 GPU, training the last layer took only 12h on the same GPU.

### 3.3 Pretraining improves predictions and reduces computation time on H3K4me3 ChIP-seq datasets across species

We then asked whether the improvement in model accuracy from pretraining is generalizable to other types of genomic datasets and to non-mammalian species. We examined ChIP-seq datasets for H3K4me3, a histone mark commonly found at promoters and enhancers [23], that were generated from rhesus macaque, mouse, and chicken cerebellum [9, 13, 26] (Figure 3A). To ensure that our findings in the ATAC-seq experiments were not a result of over-fitting on the validation set, we directly applied the hyperparameters used in the ATAC-seq experiments to experiments with H3K4me3 data without additional hyperparameter tuning.

**Figure 3.**
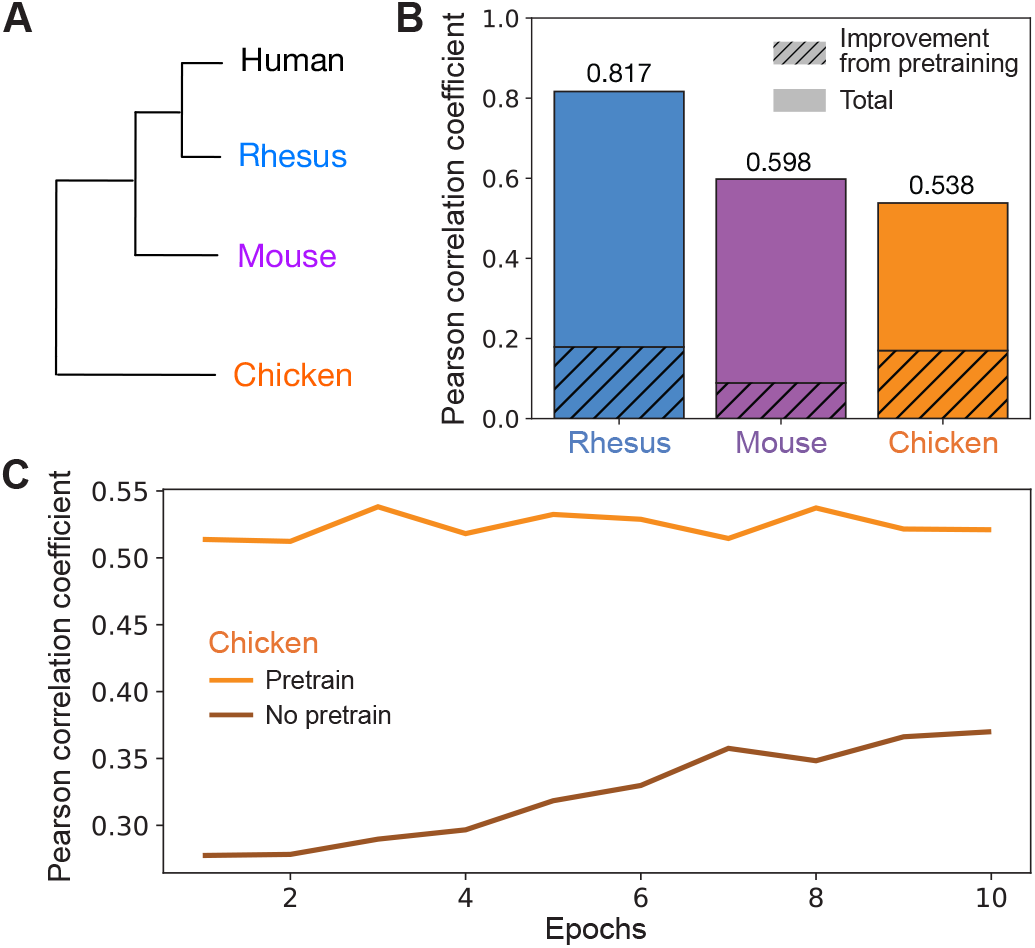
Pretraining improves performance on H3K4me3 datasets across multiple species. (A) Dendrogram of humans and species where we examined previously generated H3K4me3 ChIP-seq datasets (rhesus, mouse, chicken). (B) Validation Pearson correlation coefficients for models trained on a H3K4me3 ChIP-seq dataset from rhesus [26], mouse [13], or chicken [9]. The shaded regions are the difference between the Pearson correlation coefficient when using a model pretrained on human genomic datasets that is then fine-tuned on a H3K4me3 ChIP-seq dataset in rhesus, mouse, or chicken (“cross-species pretrained model”) and a model without pretraining. Total indicates the Pearson correlation coefficient of the cross-species pretrained model. (C) Comparison of validation Pearson correlation coefficients for the H3K4me3 ChIP-seq dataset from chicken across epochs with and without pretraining on human datasets.

Consistent with our ATAC-seq results, pretraining significantly improved model accuracy. Without pretraining, the Pearson correlation coefficient between experimental results and predictions on the test set was 0.6378, 0.5089, and 0.3684 for rhesus, mouse, and chicken respectively. With pretraining, these correlations rose to 0.8166, 0.5979, and 0.5382 for an increase of 28.03%, 17.49%, and 46.09% for rhesus, mouse, and chicken respectively (Figure 3B). Like the ATAC-seq datasets, the percent improvement in model accuracy from pretraining on human datasets was not correlated with evolutionary distance from humans. In fact, the most evolutionarily distant species (chicken) had the largest improvement in test accuracy from pretraining. This suggests that pretraining even in a distantly related species can still dramatically improve model performance upon fine-tuning in a species of interest.

Notably, pretraining also significantly reduces computation time. In all cases, for both ATAC-seq and H3K4me3 ChIP-seq datasets, we found that even after a single epoch of fine-tuning, the correlation coefficient was already higher than after 10 epochs of training from scratch (see Figure 3C for a representative plot). Thus, pretraining allows us to exceed the performance of a non-pretrained model using less than 10x the computational resources, and significantly improves model performance across diverse genomic datasets, tissues, and species.

### 3.4 Training on excessive data decreases model performance

In machine learning, training on multiple tasks can improve the model’s generalizability. However, excessive learning on diverse training tasks can impair performance on the task of interest by distributing the model’s learning ability across multiple tasks, leading to a trade-off between generalization and specialization. To determine whether there are trade-offs between generalization and specialization when training on diverse genomic datasets, we trained our simplified Enformer model on an increasing number of tracks and examined its average performance across all tracks and on a specific track of interest. We arbitrarily chose two tracks of interest: (1) H3K4me1 ChIP-seq data collected from the cerebellum of 8-week-old adult male mice and (2) DNase-seq data collected from mouse myeloid progenitor cells. We trained the Enformer model on the track of interest and 1, 50, 75, 100, 200, 500, 800, 1000, 1500, and 1642 randomly selected additional tracks. We performed three replicates for each track count, initializing the model randomly in each replicate. In all experiments, we used a learning rate of 3e-5 and trained each model for 10 epochs (see Methods).

We found that the average validation Pearson correlation coefficient across all tracks is highest when using 100-200 tracks for training (Figure 4). On our track of interest, we similarly observed the highest correlation at 100-200 tracks for training (Figure 4). When the number of tracks used during training increased beyond 200, the correlation coefficient began decreasing, indicating that excessive data is not beneficial, and may even hinder the model’s ability to learn specific tasks.

**Figure 4.**
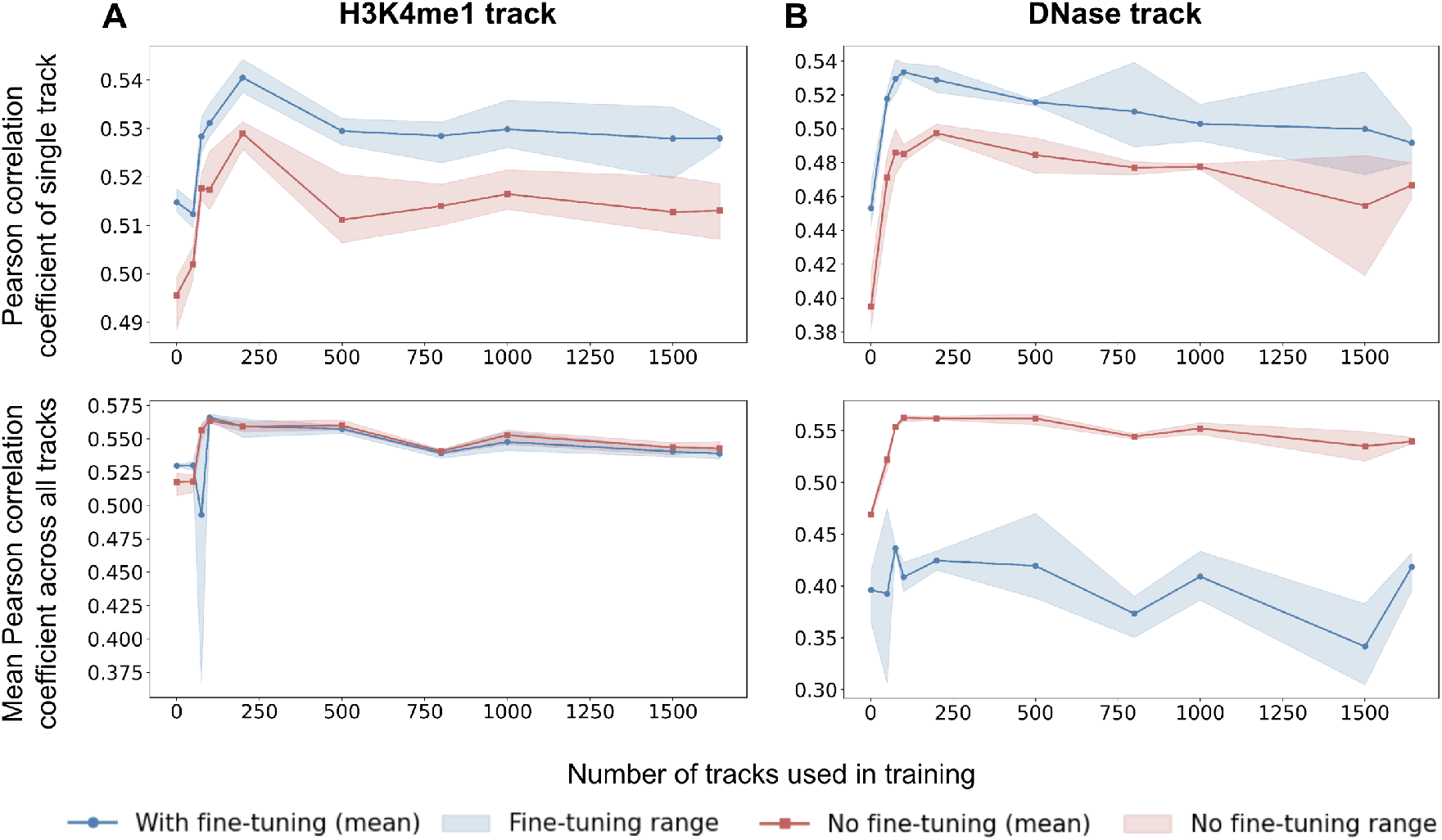
Training on an excessive number of tracks hinders model performance. (A) We trained our simplified Enformer model on a mouse cerebellum H3K4me1 track (track of interest) and 1, 50, 75, 100, 200, 500, 800, 1000, 1500, and 1642 randomly selected additional tracks for 10 epochs. Three replicates were performed. The Pearson correlation coefficient for just the H3K4me1 track (red line - top) or across all tracks (red line - bottom) is plotted. We then fine-tuned the models for another 10 epochs on just the track of interest and reported the fine-tuned Pearson correlation coefficients (blue lines). (B) We repeated the experiment in (A) using a different track of interest, a mouse myeloid progenitor DNase-seq track.

One common approach to improve a model’s specialization is through fine-tuning on specific tasks. To investigate whether a model’s specialization can be improved after training on an excessive number of tasks, we fine-tuned the models by training them for an additional 10 epochs on a single track of interest. We used a learning rate of 3e-7 for the H3K4me1 ChIP-seq track and 3e-5 for the DNase-seq track to refine the model’s performance on the target task. As expected, fine-tuning improved the Pearson correlation coefficient for the track of interest (top plots in Figure 4), while decreasing performance across all tracks (bottom plots in Figure 4). Strikingly, however, fine-tuning could not overcome the negative impact of training on an excessive number of tasks. Performance decreased when the number of tracks used in training was greater than 200 (Figure 4). These findings suggest that while fine-tuning can improve performance on a specific task, carefully calibrating the number of training tasks is critical for achieving peak performance.

## 4 Discussion

This work focuses on improving the efficiency and applicability of large deep learning models for predicting genomic datasets. We make contributions along three axes: modification of model architecture for cheaper training, using pretraining to train more effectively on single datasets, and showing that more training data does not necessarily result in more accurate models.

First, we find that deleting a linear layer from each self-attention block and a final point-wise convolutional layer from the original Enformer model improved model accuracy. We also show that the model can produce comparable results when the number of self-attention blocks is reduced from 11 to 5. While it is difficult to be sure why removing layers improved model accuracy, we hypothesize that these particular modules were only marginally useful for representing a good prediction value while significantly increasing the parameter count. Since classical statistics suggests that increased parameter count can increase training difficulty [27], this may explain the advantage of removing these layers. It remains to be determined whether these modifications would also improve the Enformer model when it is trained in the computationally intensive setting used by the original authors [5].

Although these modifications resulted in improvements over the baseline Enformer model at our smaller computational budget, this simplified model did not reach the 0.625 Pearson correlation coefficient reported by the Enformer authors [5] when trained on both human and mouse datasets. Nevertheless, we find that using these initial weights from pretraining on human datasets to then fine-tune on non-human datasets significantly improved performance over training from scratch and allowed us to reach similar or higher Pearson correlation coefficients (as high as 0.8 in many cases). This demonstrates that state-of-the-art predictions can be achieved even on a limited computational budget by tailoring the model architecture and using pretraining.

We posit that additional modifications to the model architecture may further improve performance. For instance, parameter-efficient fine-tuning strategies such as low-rank adaptation [11] are now popular and effective techniques for fine-tuning large language models in natural language processing, and could be of potential use in genomics as well. In addition, recent alternatives to attention-based architectures based on state-space models have proved effective on unsupervised DNA modeling and enhancer prediction tasks [19], suggesting that more dramatic architectural modifications may also prove effective.

Further, we show that as the number of tracks used during training increases, the correlation coefficient starts to decrease, indicating that excessive data can hinder the model’s learning ability and reduce overall performance. A similar phenomenon named the “loss of plasticity” [1, 18], where learning from an excessive number of previous tasks can affect the agent’s ability to adapt to novel tasks, is also observed in lifelong reinforcement learning (RL) [2, 28]. Our findings from fine-tuning on a single track of interest suggest that while fine-tuning helps to improve the model’s performance after training on excessive data, it does not fully mitigate the tradeoff between generalization and specialization (i.e., the model’s ability to perform well on a specific task). Thus, training on all available data followed by fine-tuning on a specific task may not be the optimal strategy for balancing both specialization and generalization.

Overall, our results strongly suggest that pretraining on human genomic datasets improves prediction accuracy and reduces computational cost and training time across different datasets, tissues, and species. This indicates that a model trained on human data appears to learn sequence representations that exploit a conserved sequence-to-function grammar across species, and opens exciting new frontiers where deep learning models in non-model organisms or in computationally constrained environments can be built using minimal experimental data and computational power. Further, our results imply that models incorporating newly generated human data do not have to be trained from-scratch but could be built cheaply by fine-tuning existing human models. The development of new models will allow us to study how sequence grammar in regulatory regions change both across species to drive evolution and also between individuals of the same species due to normal and disease-associated genetic variation. These models can also facilitate experimental efficiency, by allowing researchers to perform *in silico* genetic experiments to identify promising candidates for experimental study. Continued efforts to reduce the computational cost of these models while improving model accuracy will democratize the usability of these models in academic settings and accelerate insights into gene regulation both across and within species.

## 6 Author contributions statement

F.H., Y.W., J.H.T.S., and A.C. conceived the experiments. F.H. and Y.W. conducted the experiments. F.H., Y.W., J.H.T.S., and A.C. analyzed the results. F.H., Y.W., J.H.T.S., and A.C. wrote and revised the manuscript.

## 7 Acknowledgments

J.H.T.S. is supported by NIH/NIMH grant no. K99MH136290, and A.C. is supported by NSF grant no. 2211718.

